# Voxel-wise T_2_ relaxometry of Normal Pediatric Brain Development in 326 healthy infants and toddlers

**DOI:** 10.1101/277038

**Authors:** Vladimir S. Fonov, Ilana R. Leppert, G. Bruce Pike, D. Louis Collins, Brain Development Cooperative Group

## Abstract

Quantitative T_2_ data from an NIH-sponsored multi-center study of Normal Brain Development was used to perform automatic voxel-wise analysis of the changes in T2 evolution in the brain in healthy children within the age range from birth to 5 years. All data were non-linearly registered into a common coordinate space. The T_2_ parameters were estimated by 2 point fitting from the PD-weighted and T2-weighted image data, or by least-squares fitting of 4 data points when addition intermediate weighting images were available. The main result of this study is voxel-level map of monoexponential evolution of T_2_ in this age range indicating the delay (in months) and the rate (in 1/months) of development. The automatic maps are compared to manual region-of-interest based estimates of T_2_ evolution.

## Introduction

It is known that T1-weighted image contrast between the grey and white matter in the MRI is reversed during the first 4-6 postnatal months. The same is true for T2-weighted contrast up to 9-10 months. The actual timing of the change depends on the imaging sequence, field strength and the brain region. In this study data from an NIH-sponsored multi-center study of Normal Brain Development (NIH pediatric database (Almli, Rivkin et al.)) was used to perform automatic voxel-level analysis of the change of T2 relaxation over time for children 1-60 months old. The goal of this study was to perform voxel-level analysis of the maturation throughout the brain and quantify the differences in rate and time-delay between different anatomical regions.

## Methods

Our goal is to use data from multiple subjects at different ages to build relaxometry maps. However, the nature of the problem is complex: there are dramatic changes of the contrast between subsequent scans of the same subject over time and there are significant changes of the shape and size of the brain between subjects and over time. These issues required us to develop a special data processing technique to be able to co-register scans from the different subjects with different ages in a common coordinate system. In order to achieve these requirements we have used following approach:

- All subjects were subdivided into groups based on their age.
- Within each group, an anatomical average template was constructed using nonlinear algorithm (Fonov, Evans et al. 2011) using the T1w image modality.
- Nonlinear registration was performed between groups using Mutual Information (Mattes, Haynor et al. 2001),(Ibanez) as a cost function.
- For each scan the cumulative nonlinear mapping was created by concatenating mappings from the scan to the age-specific average template, and from the age-specific average to the 44-60 month old (mo) template.
- Resulting nonlinear mappings were used to resample individual T_2_ relaxometry maps, estimated using data from each subject-timepoint, into the common coordinate system.

### Average anatomical template

The technique of making an average anatomical template is described in (Fonov, Evans et al. 2011). The problem can be formulated as following: given a set of 3D volumes (*I*_*1*_ … *I*_*n*_), our objective is to find a 3D template *J*, such that (Equation 1), where *X*_*i*_ are individual 3D mappings of each volume *I*_*i*_ to match the template (Equation 2): under elastic body deformation constraint (Miller, Banerjee et al.). Under these assumptions it is possible to express *X*_*i*_ as a deformation vector field: 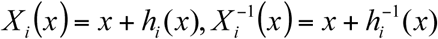, where *h(x)* may be defined at the discrete grid with given distance (step size) between nodes – as it is used in the ANIMAL algorithm

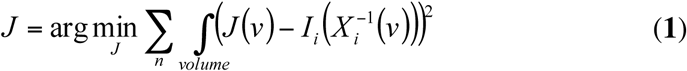

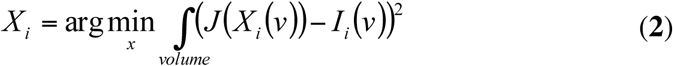

Using these ideas we have developed following algorithm:

1. Given *J*_*k*_ – the template, for each scan *I*_*i*_ calculate *X*_*i,k*_ – mappings from the template to a scan, using the 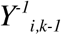 (inverse corrected mappings) from the previous iteration as a starting point (use identity for the first iteration.)
2. Calculate the mean shift of the current template (Equation 3)

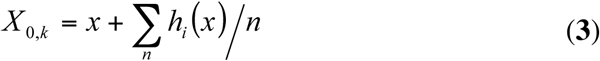
3. Calculate corrected inverse mappings: 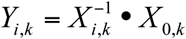
4. Apply corrected inverse mappings to individual subjects and generate an average which will be used as a new template (Equation 4):

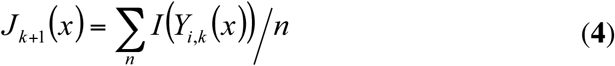
5. Repeat from step 1, until convergence is reached

The method is essentially dependant of the initial template and the possibility to find the nonlinear mapping between the given template and an individual scan. For the practical reasons we have bootstrapped the technique by doing manual linear registration of 100 scans of subjects of different ages to the MNI152 space, followed by removal of the average scaling to represent the average brain size of the population. These scans were then used to create first version of the age dependant average anatomical templates, which were used as starting points for the algorithm described above.

### T_2_ relaxometry

Each dataset was processed in the following way. All dual echo scans were registered linearly to the T1w scan using a mutual information cost function. The resulting transformation was concatenated with a nonlinear mapping from the individual scan to the age-specific template and then with the mapping to the oldest (44-60 mo) anatomical template. The concatenated registration parameters were applied to the dual-echo scans to map all information from all subjects into the common coordinate system. All images were manually inspected for the quality of registration. *T*_2_ was estimated in each dataset in a voxel-wise manner using linearized equation (*ln*(*S*_*i*_)=*ln*(*S*_0_)-*TEi/T2*) where *S*_0_ is the equilibrium signal and *S*_*i*_ is the signal is corresponding echo time *TE*_*i*_.

Regressions of *T*_2_ over age were performed in a voxel-by-voxel fashion using all images that passed QC. A mono-exponential (*T*_2_=*A+B*exp(-t*C)*) model was used. For the sake of presentation we found it is better to represent results using a different notation: *T*_2_=*A*(1+exp(-(t+D)*C))*. This method facilitates interpretation of the 3D relaxometry maps created. This way, the parameter *A*, expressed in milliseconds, can be interpreted as the asymptotic average *T*_2_ after maturation. The parameter *C,* expressed in 1/months, is the rate of change of *T*_2_ with age. Finally, the parameter *D=-ln(B/A)/C,* expressed in months, is the relative delay of the process (negative values correspond to the relative delay in maturation and positive is advance in maturation), compared to the average.

To assess the accuracy of the automatic method, a human rater manually identified several anatomical regions of interest (ROI) on the native (unprocessed) images and calculated corresponding *T*_2_. These values where then were compared to the *T*_2_ values extracted by the automatic technique, based on similar ROIs selected on the average anatomical template of 44-60 mo.

## Materials

Longitudinal MRI scans from 114 normal healthy children were acquired across 11 age cohorts (Almli, Rivkin et al.), yielding a total of 346 datasets. See Fig. 1 for the age distribution. Each dataset included one multislice T1w scan (TR=500ms TE=12ms) and a multislice dual echo PDw/T2w scan (TR=3500ms, TE=14,112ms). Some datasets included an additional acquisition of dual echo scans with longer TE=83,165ms. All calculations were performed in a Linux environment using tools from the MINC packages and ITK library (Ibanez).

**Fig. 1.**
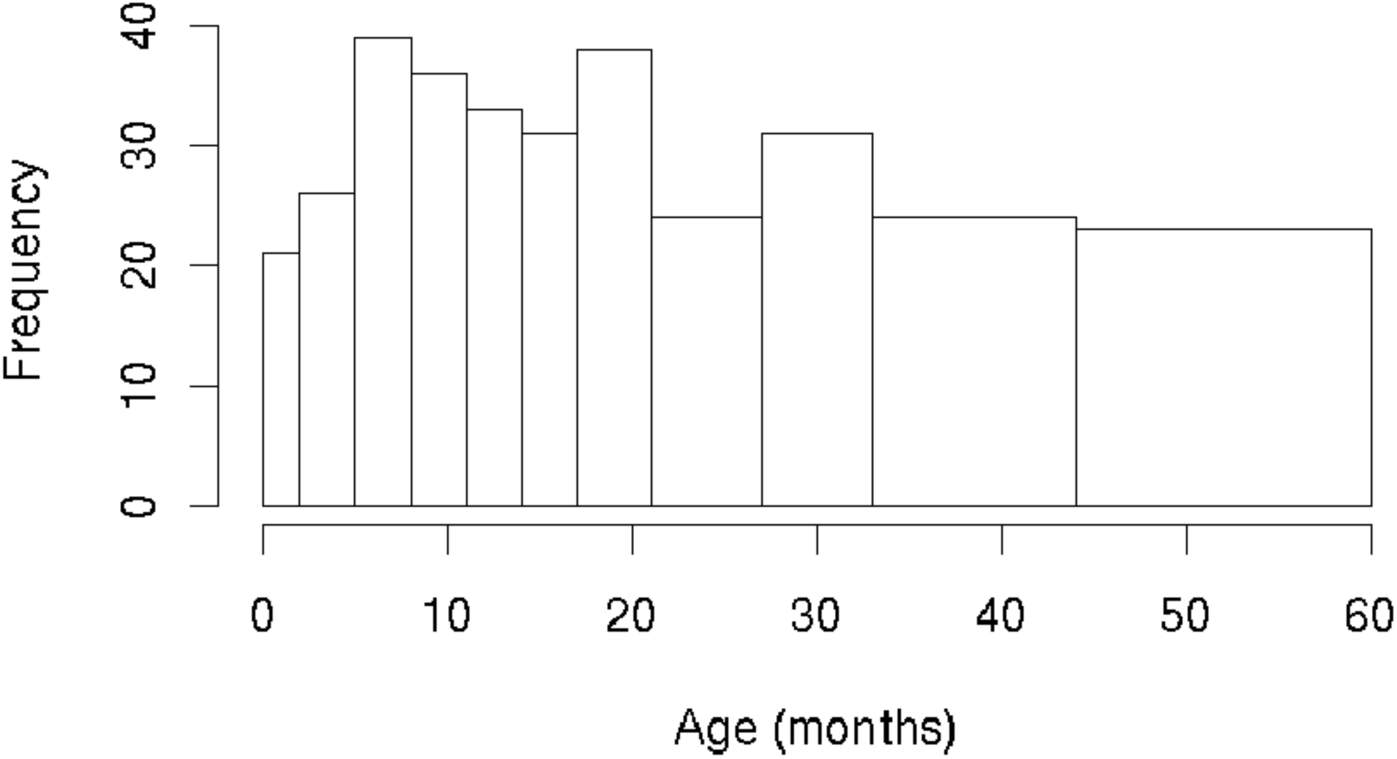
Histogram of age distribution of the datasets (n=346). Each of the 11 boxes in the histogram corresponds to one of the age ranges used to create a T1w average template (see Fig. 2).

## Results

Average anatomical templates were successfully created using 346 datasets; see Fig. 2 for the illustration on the T1w modality, templates are available at http://nist.mni.mcgill.ca.

**Fig. 2.**
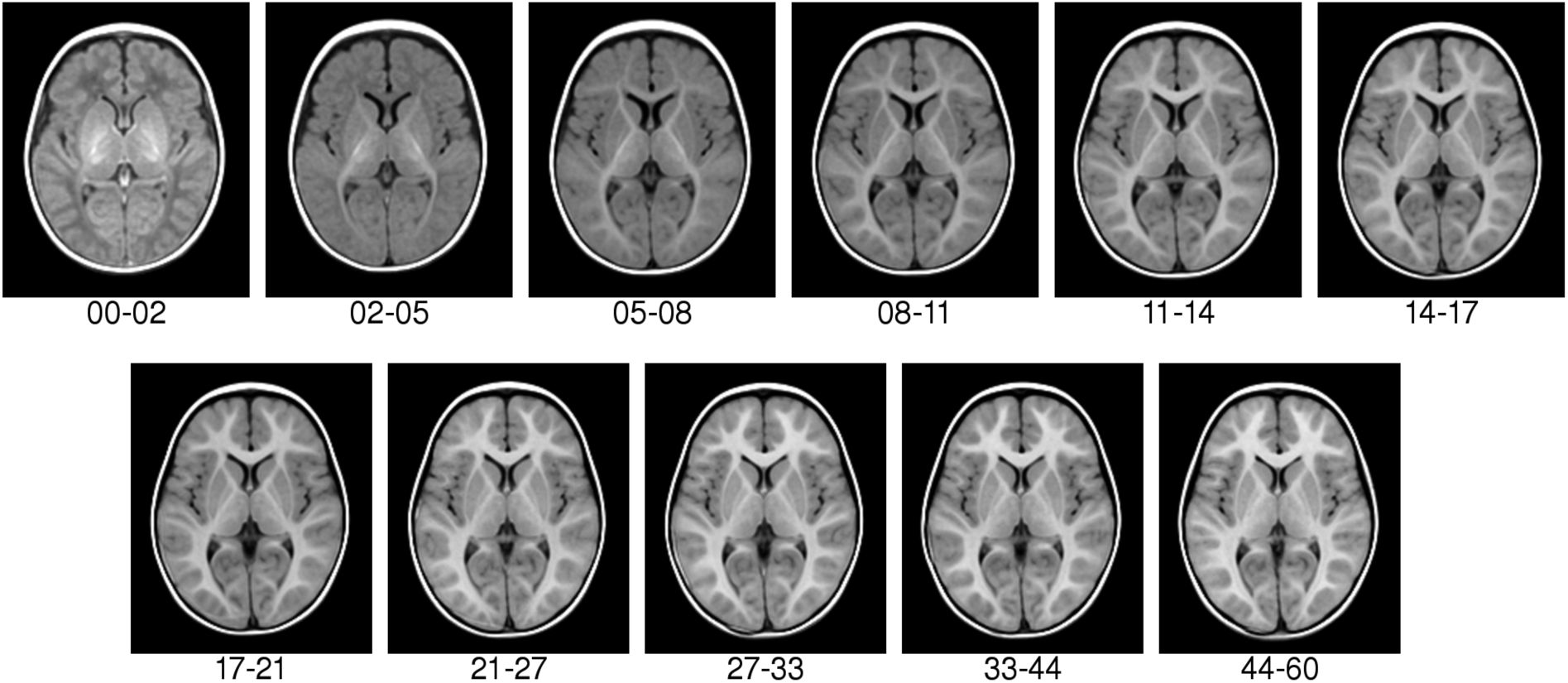
Average anatomical templates created by the iterative non-linear inter-subject registration technique (T1w modality; ages in months). One can see the change in T1w contrast with time, corresponding roughly to the myelination of the white matter with time. Note also the anatomical detail, especially at the cortex, demonstrating the high quality non-linear inter-subject registrations.

Out of 346 datasets, 326 have successfully passed all stages of automated *T2* processing, and resulting regressions show significant differences of change of *T2* between different regions of the brain during the development. Volumetric maps of A, D and C parameters and adjusted coefficient of determination R2 are shown in Fig. 3.

**Fig. 3.**
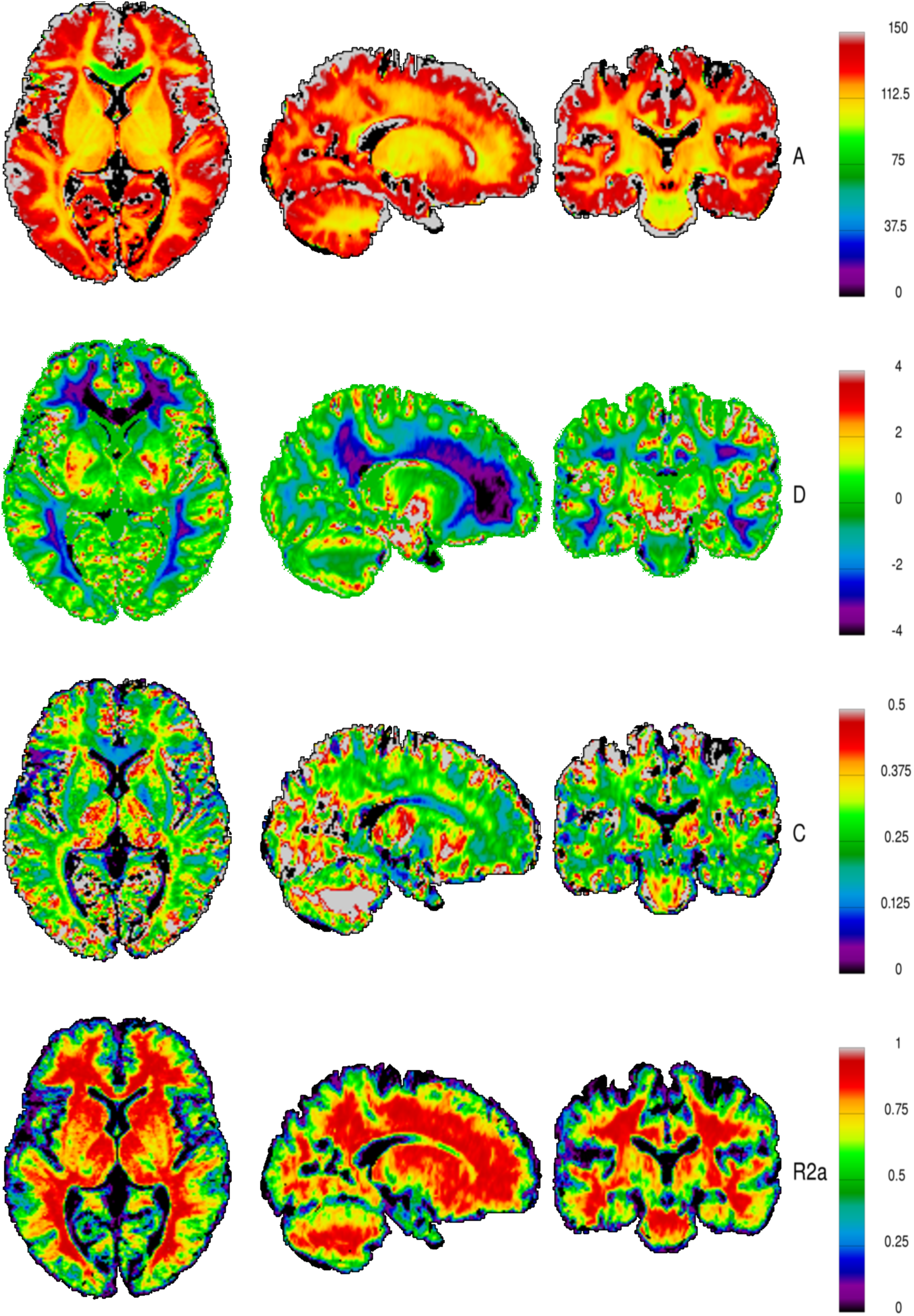
Voxel-wise regression *T*_2_=*A*(1+exp(-(t+D)*C)), A* represents asymptotic T_2_ after maturation and is expressed in milliseconds, D – the delay (when negative) or advance (when positive) of relative maturation expressed in months, C is the rate of maturation, expressed in 1/months and R2a is adjusted coefficient of determination R^2^.

Results of this study are consistent with a manual ROI-based analysis (Thalamus, Minor Forceps, Major Forceps) of T2-relaxometry performed on the same datasets in (Leppert, Almli et al.), see Fig. 4 for the comparison. The discrepancies between automatic and manual results for Major Forceps ROI for the subjects 0.5-1.0 mo may be caused by the large anatomical inter-subject variability of this area and also by the rapid maturation process happening during this age.

**Fig. 4.**
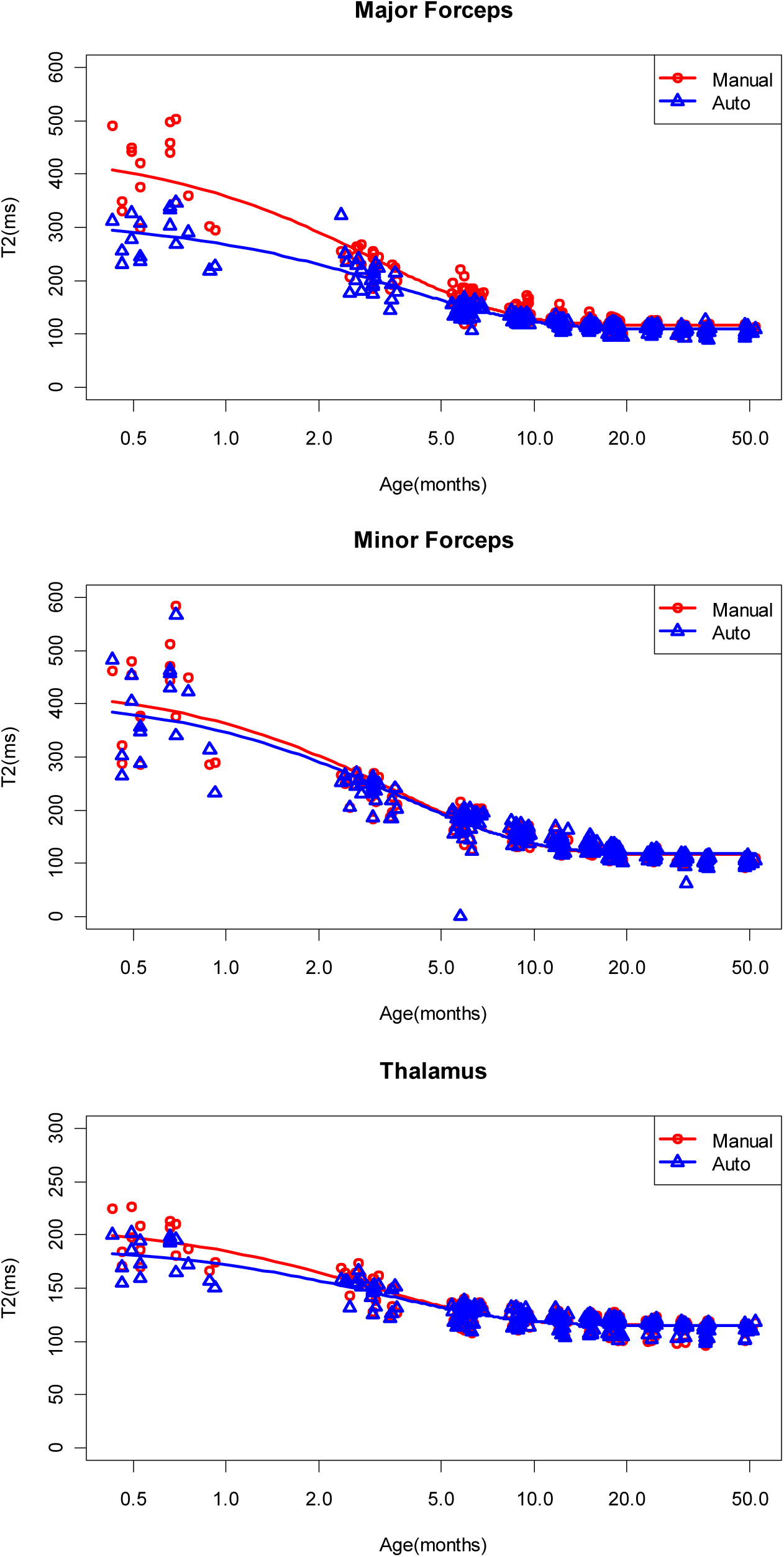
Regional T_2_ regression, comparison of manually obtained (red) and automatically calculated (blue) results estimates of T2. Note that time is shown in a logarithmic scale.

## Discussion and Conclusions

We have described an automated procedure to process T_2_ relaxometry information in a large cohort of young subjects and validated the method. Our results are consistent with the results of our manual ROI based technique with the exception of Major Forceps for subjects aged 0.5-1.0 mo. This discrepancy may be caused by the rapid maturation process occurring in this area for the given age range, corresponding with significant changes in the geometrical shape of the brain in this region, confounding our inter-subject co-registration process.

In general, our findings are consistent with the known white matter maturation pattern: from central to peripheral, from inferior to superior and from posterior to anterior (Barkovich, Kjos et al.). For example, from the sagittal view of the *C* parameter map in Fig 3, the rate of T_2_ decrease in the corpus callosum is inferior to that in more peripheral white matter. This spatial difference could reflect more advanced myelination in the central white matter, such that at birth, the structures have nearly reached full maturation and thus exhibit a slower evolution with time. The posterior to anterior progression is more visible in the *D* parameter map, where occipital white matter exhibits a more pronounced delay in reaching maturation as compared to frontal white matter. The R^2^ (coefficient of determination) map (Fig. 3) indicates that regression is able to explain 95% of inter-subject variability of T_2_ within the bulk of the white matter, and fails in the gray matter and near the edge of ventricles. This may be due to the following: the T_2_ maturation process within the gray matter cannot be explained by a mono-exponential model and the inter-subject co-registration may be poor within the cortex.

We think that our method should allow for the study of the development of the human brain in greater detail, showing quantitative differences in timing and rate of maturation of different parts of the brain.

## References

Almli, C. R., M. J. Rivkin and R. C. McKinstry (2007). “The NIH MRI study of normal brain development (Objective-2): Newborns, infants, toddlers, and preschoolers.” Neuroimage 35(1): 308–325.

Barkovich, A. J., B. O. Kjos, D. E. Jackson and D. Norman (1988). “Normal maturation of the neonatal infant brain: nMR imaging at 1.5T.” Radiology 166: 173–180.

Fonov, V., A. C. Evans, K. Botteron, C. R. Almli, R. C. McKinstry and D. L. Collins (2011). “Unbiased average age-appropriate atlases for pediatric studies.” Neuroimage 54(1): 313–327.

Ibanez, S., Ng, Cates (2005). The ITK Software Guide.

Leppert, I. R., C. R. Almli, R. C. McKinstry, R. V. Mulkern, C. Pierpaoli, M. J. Rivkin, G. B. Pike and B. D. C. Group (2009). “T2 relaxometry of normal pediatric brain development.” Journal of Magnetic Resonance Imaging 29(2): 258–267.

Mattes, D., D. R. Haynor, H. Vesselle, T. K. Lewellyn and W. Eubank (2001). Nonrigid multimodality image registration. Medical Imaging 2001, SPIE.

Miller, M., A. Banerjee, G. Christensen, S. Joshi, N. Khaneja, U. Grenander and L. Matejic (1997). “Statistical methods in computational anatomy.” Stat Methods Med Res 6(3): 267–299.

